# Suppressor mutations that make the essential transcription factor Spn1/Iws1 dispensable in *Saccharomyces cerevisiae*

**DOI:** 10.1101/2022.06.28.498004

**Authors:** Francheska López-Rivera, James Chuang, Dan Spatt, Rajaraman Gopalakrishnan, Fred Winston

## Abstract

Spn1/Iws1 is an essential eukaryotic transcription elongation factor that is conserved from yeast to humans as an integral member of the RNA polymerase II elongation complex. Several studies have shown that Spn1 functions as a histone chaperone to control transcription, RNA splicing, genome stability, and histone modifications. However, the precise role of Spn1 is not understood, and there is little understanding of why it is essential for viability. To address these issues, we have isolated eight suppressor mutations that bypass the essential requirement for Spn1 in *Saccharomyces cerevisiae*. Unexpectedly, the suppressors identify several functionally distinct complexes and activities, including the histone chaperone FACT, the histone methyltransferase Set2, the Rpd3S histone deacetylase complex, the histone acetyltransferase Rtt109, the nucleosome remodeler Chd1, and a member of the SAGA co-activator complex, Sgf73. The identification of these distinct groups suggests that there are multiple ways in which Spn1 bypass can occur, including changes in histone acetylation and alterations of other histone chaperones. Thus, Spn1 may function to overcome repressive chromatin by multiple mechanisms during transcription. Our results suggest that bypassing a subset of these functions allows viability in the absence of Spn1.

## Introduction

Eukaryotic transcription is a complicated process that requires a multitude of factors for initiation, elongation, and termination by RNA polymerase II (Schier and Taatjes 2020). Many of these transcription factors function to overcome the barriers imposed by nucleosomes. These factors include nucleosome remodeling complexes, histone modification enzymes, and histone chaperones. The importance of these activities is emphasized by their conservation throughout eukaryotes and by the finding that many of them are essential for viability. Given that there are more factors than there are perceived functions during transcription, understanding why so many vital proteins exist and how they function is an ongoing major challenge.

Our studies have focused on histone chaperones, a large and diverse set of factors that have been implicated not only in transcription, but also in DNA replication and DNA repair (hammond *et al*. 2017; warren and Shechter 2017). Histone chaperones function in a variety of ways to modulate histone-DNA interactions in an ATP-independent fashion. In several cases, the physical interactions of histone chaperones with histones and nucleosomes have been elucidated (for example, (English *et al*. 2006; Chen *et al*. 2015; Aguilar-gurrieri *et al*. 2016; Hammond *et al*. 2016; Liu *et al*. 2020; Liu *et al*. 2021)). However, the cellular roles of histone chaperones are less well understood, at least in part because of their multi-functional roles (Hammond *et al*. 2017).

An intriguing set of three histone chaperones that associate with elongating RNA polymerase II (RNAPII) are Spn1/Iws1, Spt6, and FACT. Each of these histone chaperones is highly conserved from yeast to human, and each is essential for viability in most organisms where this has been assessed. Spt6 and Spn1/Iws1 directly interact with each other (Diebold *et al*. 2010; Mcdonald *et al*. 2010) and all three chaperones are functionally connected based on substantial genetic and molecular evidence (Duina 2011; Jeronimo *et al*. 2015; Mccullough *et al*. 2015; Pathak *et al*. 2018; Jeronimo *et al*. 2019; Viktorovskaya *et al*. 2021). However, the precise roles of each factor, the nature of their functional and physical interactions, and why they are essential for viability are not fully understood.

In this work, we have studied one of these histone chaperones, Spn1 (named Iws1 in organisms other than *Saccharomyces cerevisiae* (*S. cerevisiae*)). Spn1 was initially identified in *S. cerevisiae* (Fischbeck *et al*. 2002; Krogan *et al*. 2002; Lindstrom *et al*. 2003), where it was subsequently shown to directly interact with Spt6 (Diebold *et al*. 2010; Mcdonald *et al*. 2010). Spn1 functions as a histone chaperone based on several lines of evidence. First, *in vitro*, purified Spn1 interacts directly with histones, nucleosomes, and nucleosomal arrays, and it has modest nucleosome assembly activity (Li *et al*. 2018; Li *et al*. 2022). Second, *in vivo*, a temperature-sensitive *spn1* mutation, *spn1-K192N* (Fischbeck *et al*. 2002), causes changes in chromatin structure, resulting in altered spacing between nucleosomes and increased nucleosome fuzziness (Viktorovskaya *et al*. 2021). Third, *spn1-K192N* is suppressed by mutations that alter histone H2A (Mccullough *et al*. 2015) or that impair other chromatin-related factors, including FACT, Swi/Snf, Set2, and Rpd3S (Zhang *et al*. 2008; Lee *et al*. 2018b; Viktorovskaya *et al*. 2021). Finally, *spn1* mutations cause double mutant sickness or lethality when combined with mutations that impair other histone chaperones (Li *et al*. 2018) or histone H3K4 methylation (Lee *et al*. 2018a). Together, this evidence shows that Spn1 functions to control chromatin structure by direct interaction with histone proteins.

Spn1 associates with chromatin along coding regions, suggesting that its main function is during transcription elongation (Mayer *et al*. 2010; Reim *et al*. 2020). Studies in mammalian cells have shown that Spn1 interacts directly with several elongation factors in addition to Spt6, leading to the hypothesis that it acts as a hub for the RNA polymerase II elongation complex (Ling *et al*. 2006; Liu *et al*. 2007; Cermakova *et al*. 2021). Consistent with this idea, recent studies showed that Spn1 is required for normal transcript levels in both yeast and mammalian cells (Reim *et al*. 2020; Cermakova *et al*. 2021), as well as for mRNA splicing and histone modifications (Yoh *et al*. 2007; Yoh *et al*. 2008; Reim *et al*. 2020). Furthermore, in mammalian cells, Spn1 is essential for embryonic development and cell proliferation (Liu *et al*. 2007; Orlacchio *et al*. 2018; Oqani *et al*. 2019), and it has been implicated in cancer (Davoli *et al*. 2013; Sanidas *et al*. 2014). Thus, Spn1 plays critical roles in gene expression, development, and disease.

Given the importance of Spn1 throughout eukaryotes and its implication in human disease, we wanted to understand more about Spn1 function and why Spn1 is essential for viability. To do this, we isolated bypass mutations that suppress the inviability caused by a deletion of the *SPN1* gene. This type of approach has been highly informative in several past studies, leading to insights into how an organism can compensate for loss of an essential gene (for example, (Zhao *et al*. 1998; Biswas *et al*. 2008; Rolef ben-shahar *et al*. 2008; Torres-machorro and Pillus 2014; Chang *et al*. 2018), for a review, see (Du 2020)). In addition to single-gene studies of bypass suppressors, other studies have systematically screened for suppressors of essential genes in either *S. cerevisiae* or *Schizosaccharomyces pombe* (*S. pombe*) (Chen *et al*. 2016; Li *et al*. 2019; Takeda *et al*. 2019; Van leeuwen *et al*. 2020). The results of these studies have led to interesting ideas concerning the evolvability and the relative importance of essential genes, based on whether or not they could be suppressed. However, those surveys likely underestimated the number of essential genes that can be bypassed, as no suppressors of *spn1Δ* were identified in either of the *S. cerevisiae* studies.

Our results have shown that Spn1 can be bypassed by single mutations in at least eight different genes that control transcription and chromatin structure. In this work, we have followed up on two distinct and apparently opposite classes. Loss of one class, Set2/Rpd3S, causes increased levels of histone acetylation and we show that increased acetylation is indeed the mechanism by which loss of Set2/Rpd3 bypasses Spn1. In stark contrast, loss of the second class, Rtt109, causes decreased levels of histone acetylation. Our analysis of suppression by loss of Rtt109 suggests that the sites of acetylation are critical for their effect on Spn1 bypass. The identification of such distinct classes of Spn1 bypass suppressors suggests that Spn1 is essential due to more than one activity, and suppression of a subset of these activities allows viability.

## Methods

### Yeast strains and growth media

The *S. cerevisiae* strains with an FY name (Supplementary Table 1) were in an isogenic background derived from S288C (Winston *et al*. 1995) and were constructed by standard procedures using yeast transformation and crosses. CRISPR/Cas9 directed changes were done as previously described (Laughery *et al*. 2015). Strains that contain the *hht1-K56R* and *hht2-K56R* mutations (Mccullough *et al*. 2019), kindly provided by Dr. Tim Formosa, were derived from a cross of strain FY3462 (S288C background) by strain DY16302 (W303 background). Primers used for PCR are listed in Supplementary Table 2. Genotypes were confirmed by growth on selective media, PCR, or DNA sequencing. Strains were grown in the following liquid or solid media: YPD (1% yeast extract, 2% peptone and 2% glucose); solid YPD with G418 (200 μg/ml), hygromycin (7.5 μg/plate), nourseothricin (3 μg/plate), dimethyl sulfoxide (DMSO, 0.1%), or 1-naphthaleneacetic acid (NAA, 500 μM); synthetic complete (SC) media lacking the amino acid of interest (0.2% dropout mix, 0.145% yeast nitrogen base without ammonium sulfate and amino acids, 0.5% ammonium sulfate, 2% glucose); or SC media with 0.005% uracil and with 5-fluoroorotic acid (5-FOA, 1 mg/ml). *S. pombe* was grown in YES liquid or solid media (0.5% yeast extract, 3% glucose, 225 mg/L each of adenine, histidine, leucine, uracil, and lysine) at 32°C. Spot tests to assay growth on different media used ten-fold serial dilutions of each strain with the most concentrated spot having an OD_600_ “ #$

### Construction of strains to select for suppressors of *spn1Δ*

To allow for selection of *spn1Δ* suppressors, *S. cerevisiae* strains were built using the following steps. First, *MAT***a** and *MATα* wild-type strains were transformed with plasmid FB2701, which contains copies of the *SPN1, URA3*, and *HIS3* genes. This plasmid was generated by cloning a copy of the *URA3* gene into plasmid pCR311, kindly provided by Dr. Laurie Stargell (Fischbeck *et al*. 2002), composed of pRS313 (Sikorski and Hieter 1989) with *2xmyc-SPN1*, including 403 and 116 base pairs (bp) of upstream and downstream genomic sequences, respectively. Second, the genomic *SPN1* open reading frame (ORF) was replaced in the transformed strains with one of the following drug-resistance cassettes: *kanMX, natMX*, or *hphMX*, to obtain the *spn1Δ* parental strains listed in Supplementary Table 1.

### Selection and screen to isolate *spn1Δ* suppressors

About 50 single colonies each from strains FY3007 and FY3008 were spread as patches that occupied a quarter of a solid YPD plate. After one day of incubation at 30°C, the patches were replica plated on medium containing 5-fluoroorotic acid (5-FOA) to detect cells that had lost the *SPN1* plasmid. A subset of the plates was treated with 5000 μJ/cm^2^ of UV light using the Stratagene UV Stratalinker 2400, while the rest of the plates were left untreated. UV-induced or spontaneous suppressors were selected by incubating the 5-FOA plates at 30°C for 3 days, and then replica plating them a second time on 5-FOA. This second set of 5-FOA plates, which had reduced background growth compared to the first set, was incubated at 30°C for 5-7 days. Colonies that grow on 5-FOA could be either new mutations that suppress *spn1Δ* inviability, thus allowing plasmid loss, or could be mutations in the plasmid-borne *URA3* gene. To distinguish between these two possibilities, the 5-FOA resistant candidates were screened by replica plating for those that were also His^-^, which would indicate plasmid loss. A maximum of two 5-FOA resistant His-candidates per patch were single-colony purified on YPD. Only one candidate from each patch that had a deletion of *SPN1* and absence of *URA3* and *HIS3* was considered an independent *spn1Δ* suppressor. In exceptional cases, two candidates isolated from the same patch were considered independent if the strains showed different growth patterns on SC-lys or on YPD media at 30°C or 37°C. From this we isolated 105 mutants.

### Dominance tests

Each of the 105 *spn1Δ* suppressor strains with unidentified suppressor mutations was grown as parallel stripes on solid YPD, with a maximum of eight strains per 10-cm plate. The *spn1Δ* parental strains FY3007 and FY3008, which contain the *SPN1 URA3 HIS3* plasmid, were also grown as parallel stripes on a different set of plates. The haploid *spn1Δ* suppressor strains were mated with haploid *spn1Δ* parental strains of the opposite mating type by replica plating both sets of strains on YPD, forming a grid of perpendicular stripes, thus allowing diploids to form where the stripes intersected. As a positive control, diploids with the *SPN1* plasmid were selected by replica plating onto SC-leu -trp. Dominance was assayed by selecting for diploids without the *SPN1* plasmid on SC-leu -trp + 5-FOA. Growth on these plates suggested that the diploid strain had a dominant *spn1Δ* suppressor, while failure to grow suggested that the diploid strain had a recessive suppressor.

### Whole-genome sequencing analysis

To identify candidate suppressor mutations by WGS, we selected the 30 *spn1Δ* suppressor strains that were isolated spontaneously and that had the strongest *spn1Δ* suppression phenotype based on growth in the absence of the *SPN1* plasmid at 30°C. We grew 25 ml liquid YPD cultures for each suppressor strain, as well as the *spn1Δ* parental strains (FY3007, FY3008, FY3038, and FY3040) until saturation was reached. This took multiple days for some strains. The cell concentration was determined using a hemacytometer, and the volume of saturated culture required to obtain 4-8×10^8^ total cells was calculated. This volume of cells was pelleted, washed with 1 ml of distilled water, pelleted again, and flash frozen. Extraction and shearing of genomic DNA, library construction, and whole-genome sequencing were performed as described (GOPALAKRISHNAN AND WINSTON 2019). Although we found suppressor mutations in multiple classes of genes, we focused on the mutations that targeted the coding regions of transcription- and chromatin-associated genes for follow-up studies. Additional analyses to identify the suppressors and to check for absence of the *SPN1, URA3* and *HIS3* genes were performed using Geneious version 11.0.5 (KEARSE *et al*. 2012).

### Gene replacements to test candidate *spn1Δ* suppressors

To test the candidate *spn1Δ* suppressors in non-essential genes, we replaced the wild-type gene with either the *kanMX* or the *hphMX* drug-resistance cassette (Bergkessel *et al*. 2013), using the primers listed in Supplementary Table 2. In each case, the PCR products were column-or gel-purified and transformed into a *SPN1/spn1Δ* heterozygous diploid strain containing the *SPN1 URA3 HIS3* plasmid, which was constructed by crossing FY3007 and FY132. Transformants were selected on media containing G418 or hygromycin, and the gene replacements were confirmed by PCR. After sporulation and tetrad analysis, the haploid strain that had deletion of both *SPN1* and the candidate non-essential gene and that also contained the *SPN1 URA3 HIS3* plasmid were tested for suppression on media containing 5-FOA. To test the candidate *spn1Δ* mutations in the essential genes *POB3* and *SPT16*, we performed two-step gene replacement (Gray *et al*. 2005). A *SPN1/spn1Δ* heterozygous diploid strain was constructed by mating FY3007 and FY1856 and selecting for diploids that had lost the *SPN1 URA3 HIS3* plasmid on 5-FOA. The diploid was transformed with PCR products that had the *URA3* ORF flanked by sequences homologous to the gene of interest. Transformants were selected on SC-ura and re-transformed with PCR products that contained the candidate suppressor. Transformants that lost the *URA3* gene were selected on media containing 5-FOA. Presence of the candidate suppressor was checked by PCR and/or sequencing. Strains that had deletion of *SPN1*, the candidate suppressor, and the *SPN1 URA3 HIS3* plasmid were tested for suppression on media containing 5-FOA.

### Plasmid complementation to test *spn1Δ* suppression by alteration of *TFG1*

To test suppression by alteration of *TFG1*, we performed the following steps. First, we replaced the *TRP1* ORF with the *hphMX* cassette in both the FY3531 suppressor strain, which has *spn1Δ* and a candidate suppressor in *TFG1*, and the FY3463 wild-type strain. We then transformed both strains with plasmid pGH262 (Lindstrom *et al*. 2003), kindly provided by Dr. Grant Hartzog, which contains *URA3* and an untagged version of *SPN1*, including 479 bp of upstream and downstream genomic sequences. The transformed strains were then transformed a second time with one of two *TRP1*-containing plasmids: (1) an empty pRS314 plasmid (Sikorski and Hieter 1989) or (2) pRS314-TFG1, kindly provided by Dr. Stephen Buratowski. The four resulting strains, FY3464-FY3467, were tested for *spn1Δ* suppression on SC-trp containing 5-FOA. Inviability on 5-FOA of the FY3531 *spn1Δ* suppressor strain with pRS314-TFG1 would suggest that alteration of *TFG1* is responsible for *spn1Δ* suppression.

### Chromatin immunoprecipitation-sequencing (ChIP-seq)

Spn1 degron strains FY3477 and FY3478 were grown in triplicate as 170 ml YPD cultures at 30°C to OD_600_ ≈ 0.6. The *S. pombe* strain FWP10 was grown in YES at 32°C until an OD_600_ ≈ 0.6. The FY3477 and FY3478 cultures were diluted two-fold with YPD and divided into two equal volumes with OD_600_ ≈ 0.3. For Spn1 depletion, one of the cultures from each train was treated with 25 μM indol-3-acetic acid (IAA) auxin (dissolved in DMSO) for 90 minutes, and for non-depletion, DMSO was added to the other culture as previously described (Reim *et al*. 2020). To assay Spn1 depletion, 10 ml of each of the *S. cerevisiae* cultures were collected by centrifugation and whole cell lysates (Matsuo *et al*. 2006) were prepared to measure Spn1 levels by Western blot as in (Reim *et al*. 2020), using a 1:6,000 dilution of anti-Spn1 antisera (kindly provided by Dr. Laurie Stargell) and a 1:10,000 dilution of anti-actin (Abcam, ab8224) as a loading control. In addition, 150 ml of each culture were transferred to a flask that was at room temperature, and 36 ml of 4°C YPD were added to each to reach room temperature. These cultures were then fixed with 1% formaldehyde for 30 minutes while shaking at room temperature. The crosslinking was quenched by adding glycine to each culture to a final concentration of 125 mM and by shaking for 10 minutes at room temperature. The cells were pelleted, washed twice with cold 1x TBS (100 mM Tris, 150 mM NaCl pH 7.5), and flash frozen. FY3477 and FY3478 cell pellets were resuspended in 800 μl of LB140 (50 mM HEPES-KOH pH 7.5, 140 mM NaCl, 1 mM EDTA, 1%. Triton X-100, 0.1% sodium deoxycholate, 0.1% SDS, 100 μg of leupeptin, 100 μg of pepstatin A, 8.71 mg of PMSF, 3.08 mg of 1M DTT), and lysed by bead-beating at 4°C for 8 minutes with 4-minute incubations on ice after every minute of bead-beating. The *S. pombe* FWP10 pellets were treated similarly, but they were lysed by bead-beating at 4°C for 11 minutes. The lysate was collected by centrifuging the samples at 14,000 rpm for 5 minutes at 4°C. After discarding the supernatant, the pellet was washed with LB140. The final pellet was washed in 580 μl of LB140 and sonicated in a QSonica Q800R machine for 10 minutes (30 seconds on, 30 seconds off, 70% amplitude) to obtain fragments of about 100-500 bp. The protein concentrations of all samples were measured by Bradford (Bradford 1976). To prepare the IP reactions, about 250 μg of *S. cerevisiae* chromatin was used as starting material, and 10% w/w *S. pombe* chromatin was added to each sample as a spike-in control (Orlando *et al*. 2014; Gopalakrishnan *et al*. 2019). Samples were brought to 250 μl in LB140 buffer, then 250 μl WB140 buffer (50 mM HEPES-KOH pH 7.5, 140 mM NaCl, 1 mM EDTA, 1% Triton X-100, 0.1% sodium deoxycholate) was added. Then, 40 μl of each sample were set aside as input for later processing at the reverse crosslinking step, described later. Two sets of IPs were prepared per sample with the following antisera: 3 μl of anti-H4 (AV94), kindly provided by Dr. Alain Verreault, and 16.7 μl of anti-H4ac (Millipore Sigma, 06-866, which targets histone H4ac on lysines 5, 8, 12, and 16), volumes that were optimized by ChIP-qPCR. The immunoprecipitation reactions were incubated at 4°C with end-over-end rotation overnight (∼14-16 hours). Aliquots of Dynabeads Protein G for Immunoprecipitation (Invitrogen) and Dynabeads Protein A for Immunoprecipitation (Invitrogen) were transferred to Eppendorf tubes, and the buffer was removed. A volume of WB140 equivalent to the initial volume of beads was used to wash the beads twice with WB140, and the beads were kept in the second wash to make a slurry. Fifty μl of the Protein G slurry were added to each of the H4 IPs, 278 μl of the Protein A slurry were added to the H4ac IPs, and the reactions were incubated at 4°C with end-over-end rotation for four hours. The beads were washed twice (first wash for one minute and second wash for five minutes with nutation) with WB140, twice with WB500 (50 mM HEPES-KOH pH 7.5, 500 mM NaCl, 1 mM EDTA, 1% Triton X-100, 0.1% sodium deoxycholate), twice with WBLiCl (10 mM Tis-HCl pH 7.5, 250 mM LiCl, 1 mM EDTA, 0.5% NP-40, 0.5% sodium deoxycholate). The beads were then washed once with TE (10 mM Tris-HCl pH 7.4, 1 mM EDTA) for 5 minutes. The IP material was eluted from the beads twice with 100 μl of TES (50 mM Tris-HCl pH 7.4, 10mM EDTA, 1% SDS) at 65°C for 30 minutes first and then for 15 minutes. The saved inputs were thawed, and 60 μl of LB140 and 100 μl of TES were added. To reverse crosslinking, the eluates and the inputs were incubated at 65°C overnight. To degrade RNA, 5 μl of RNAse A/T1 (Thermo Fisher, EN0551) were added to each sample, and the samples were incubated at 37°C for one hour. To degrade protein, a mix of 7 μl of proteinase K (Roche), 1 μl of glycogen (Sigma G1767-1VL, 20 mg/ml), and 250 μl of TE were added to each sample, and the samples were incubated at 50°C for two hours. A DCC-5 kit (Zymo Research, D4014) was used to purify DNA (20-μl elution). The libraries were built with 10.25 μl of DNA as starting material and using the GeneRead DNA Library I Core Kit (Qiagen, half of the volumes suggested by the manufacturer instructions were used) and custom barcodes (WONG *et al*. 2013). The samples were purified twice using 0.7x volume SPRI beads (A63881). PCRs were performed to determine number of amplification cycles, which ranged from 11 to 19, and to perform final library amplifications. The size distribution, quality, and concentration of the samples was assessed by running them in an Agilent Bioanalyzer. The resulting 45 ChIP-seq libraries were sequenced using the Illumina NextSeq High 75 cycles platform at the Harvard Bauer Core Facility. The data were analyzed as in (Reim *et al*. 2020)

## Results

### Isolation of mutations that suppress the inviability caused by deletion of the *SPN1* gene

To isolate suppressors of a deletion of the essential *SPN1* gene *(spn1Δ)*, we followed a two-step selection and screening protocol that identified viable *spn1Δ* strains (Figure 1A, Methods). We first isolated 105 independent suppressor candidates; of these 104 were recessive and one was dominant. To identify the causative mutations, we performed whole-genome sequencing of the 30 candidates with the strongest suppression phenotypes (29 recessive, one dominant). Sequencing of the mutants, all of which were spontaneous isolates, identified 7-21 mutations per genome (Supplementary Table 3). We then tested candidate genes by gene replacement analysis, focusing on genes known to be involved in transcription and chromatin structure. This analysis led to the identification of suppressor genes in 27 of the 30 strains (Supplementary Table 4). The mutations from the 27 strains identified at least eight genes that suppress *spn1Δ* to different degrees (Figure 1A, 1B, Supplementary Table 4).

**Figure 1.**
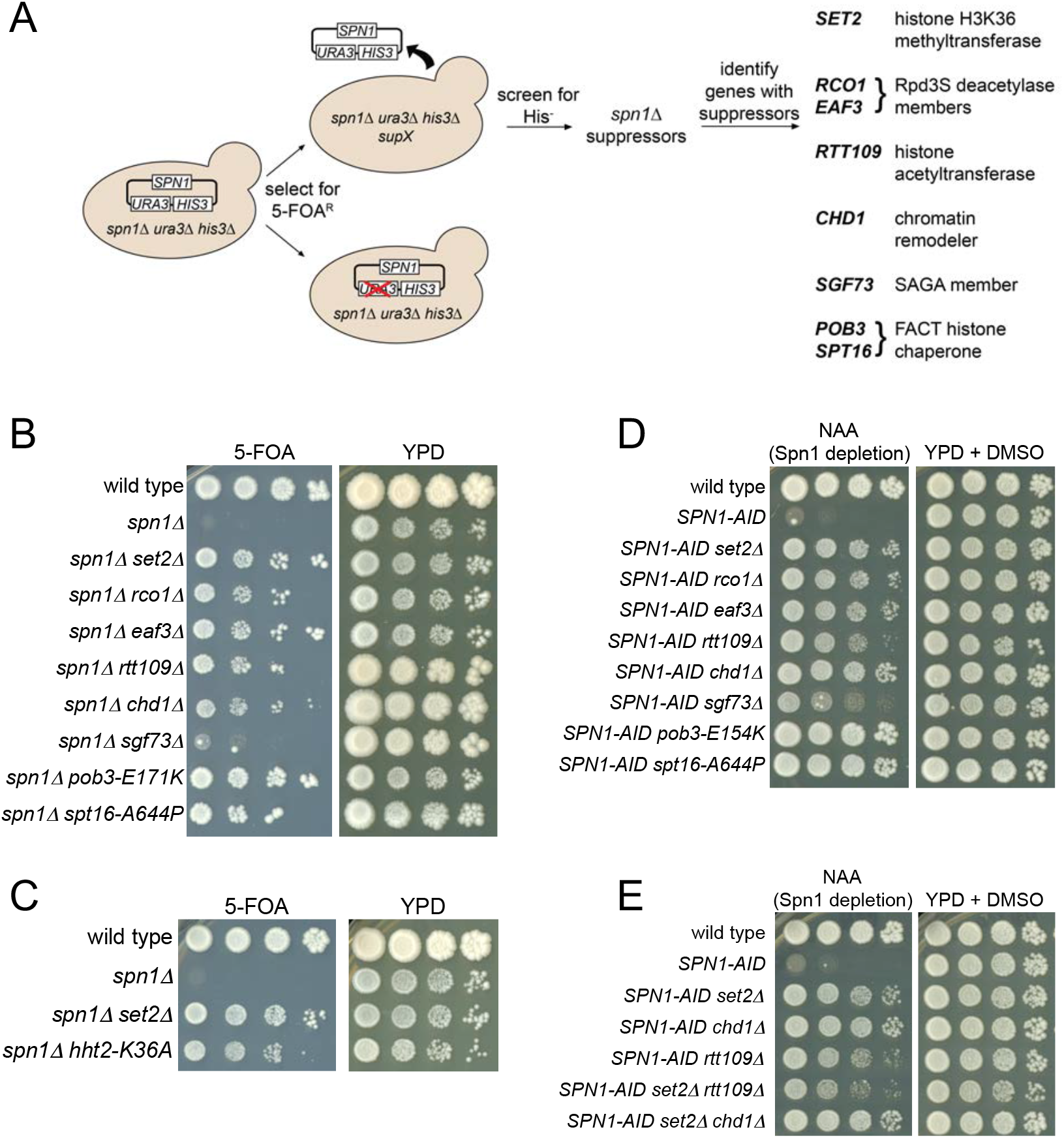
The isolation and analysis of Spn1 bypass suppressors. (A) A schematic showing the selection and subsequent screen used to isolate Spn1 bypass suppressors. Briefly, *spn1Δ* yeast strains that also have deletions of *HIS3* and *URA3* were grown. These strains are viable because they contain a wild-type copy of the *SPN1* gene in a plasmid, which also has wild-type copies of *URA3* and *HIS3*. Suppressors were selected on medium containing 5-fluoroorotic acid (5-FOA), which counterselects for cells that have *URA3*. To screen for the suppressor strains that lost the plasmid, rather than those that acquired a *ura3* mutation, the 5-FOA resistant candidates were screened for those that were also His-. We isolated 105 independent *spn1Δ* suppressors. Whole-genome sequencing of the 31 strongest mutants, followed by gene replacements, identified eight genes with *spn1Δ* suppressors. (B) Dilution spot tests of *spn1Δ* suppressors. Shown are 5-FOA (suppression) and YPD (permissive) plates after seven days at 30°C. (C) Suppression of *spn1Δ* by *set2Δ* and by H3K36A. Plates are shown after five days at 30°C. (D) The suppressors of *spn1Δ* also suppress Spn1 depletion mediated by *SPN1-AID*. Plates are shown after three days at 30°C. Plates labeled NAA contain 500 μM 1-naphthaleneacetic acid to enable Spn1 depletion. Here we tested suppression of *pob3-E154K*, while in panel A we tested *pob3-E171K*. Both mutations target the dimerization domain of Pob3 as previously described (Viktorovskaya *et al*. 2021) (E) Analysis of suppression by double mutants, shown after three days of incubation at 30°C.

The eight genes encode proteins that function in different aspects of transcription and chromatin structure. Two of the genes, *SPT16* and *POB3*, encode the two subunits of the essential FACT histone chaperone complex. Some of the *spt16* and *pob3* suppressor mutations identified here were recently described (Viktorovskaya *et al*. 2021). The other six genes encode non-essential proteins: Set2 is the sole *S. cerevisiae* H3K36 methyltransferase (Strahl *et al*. 2002; Keogh *et al*. 2005; Kizer *et al*. 2005; Pokholok *et al*. 2005); Rco1 and Eaf3 are subunits of the Rpd3S histone deacetylase complex, whose deacetylase activity on histones H3 and H4 requires H3K36 methylation by Set2 (Carrozza *et al*. 2005; Joshi and Struhl 2005; Keogh *et al*. 2005; Li *et al*. 2007a; LI *et al*. 2007b; Drouin *et al*. 2010; Govind *et al*. 2010); Chd1 is an ATP-dependent chromatin remodeler (Woodage *et al*. 1997; Tran *et al*. 2000); Rtt109 is a histone acetyltransferase with multiple targets (Schneider *et al*. 2006; Driscoll *et al*. 2007; Han *et al*. 2007; Tsubota *et al*. 2007; Fillingham *et al*. 2008; Albaugh *et al*. 2011; Abshiru *et al*. 2013; KUO *et al*. 2015; Cote *et al*. 2019) and Sgf73 is a member of the SAGA coactivator complex, required for anchoring the SAGA histone deubiquitylation module (Helmlinger *et al*. 2004; Mcmahon *et al*. 2005; Lee *et al*. 2009; Yan and Wolberger 2015). For all six non-essential genes, we found that a complete deletion could suppress *spn1Δ* to varying degrees (Figure 1B), demonstrating that, for each of these six genes, suppression is caused by loss of function.

The suppression of *spn1Δ* by *set2Δ* suggested that loss of histone H3K36 methylation suppressed *spn1Δ*. However, Set2 may have other targets in yeast in addition to histone H3K36, as has been shown for its mammalian counterpart, SETD2 (Park *et al*. 2016). To test whether it is specifically the loss of H3K36 methylation that is responsible for *spn1Δ* suppression, we constructed strains that produced only a mutant histone H3, H3K36A, which cannot be methylated at position 36. Our results showed that this mutation suppressed *spn1Δ* nearly as efficiently as did *set2Δ* (Figure 1C). Therefore, we conclude that loss of H3K36 methylation is responsible for *spn1Δ* suppression in a *set2Δ* strain.

To facilitate the analysis of Spn1 bypass suppressors, we switched from using *spn1Δ* to using strains in which we could conditionally deplete Spn1 using an auxin-inducible degron fused to the *SPN1* gene (Nishimura *et al*. 2009; Reim *et al*. 2020). Our previous work showed that we could rapidly deplete cells of Spn1 protein as judged by both Western analysis and chromatin immunoprecipitation (Reim *et al*. 2020). Using this system, we re-analyzed the eight suppressor mutations by their ability to grow on plates that contain the auxin 1-naphthaleneacetic acid (NAA; see Methods). Our results showed that all eight mutations suppressed the growth defect of *SPN1-AID* on NAA plates (Figure 1D), thus validating this as a useful system for studying Spn1 bypass suppressors.

Using this Spn1 depletion system, we tested whether combinations of suppressor mutations would confer stronger suppression than single suppressor mutations. This was of interest as several of our original isolates, although spontaneous, contained mutations in multiple suppressor genes (Supplementary Table 4). We constructed two double mutant combinations, *set2Δ chd1Δ* and *set2Δ rtt109Δ*. These choices were made because of evidence, presented later, that *set2Δ* bypasses Spn1 by a distinct mechanism from either *chd1Δ* or *rtt109Δ*. In addition, *set2 chd1* double mutants were found in our original suppressor strains. Our results showed that suppression was not detectably greater in the double mutants than in any of the single mutants (Figure 1E). Presumably, the *set2* and *chd1* mutations in the original isolates were weaker mutations than the deletions, such that combining them would result in greater suppression than by either mutation alone.

### Bypass of Spn1 by *set2Δ* occurs via elevated histone acetylation levels

We suspected that Spn1 bypass by *set2Δ, rco1Δ*, and *eaf3Δ* might occur via increased histone acetylation, as several previous studies showed that histone H3 and H4 acetylation levels are elevated across coding regions in those mutants (Vogelauer *et al*. 2000; Carrozza *et al*. 2005; Joshi and Struhl 2005; Keogh *et al*. 2005; Li *et al*. 2007b; Li *et al*. 2009; Drouin *et al*. 2010; Govind *et al*. 2010; Venkatesh *et al*. 2012). We considered that suppression by elevated histone acetylation levels could be due to two possible defects caused by Spn1 depletion. First, Spn1 depletion might cause decreased levels or altered localization of histone acetylation. Given the essential nature of histone acetylation by the histone H4 acetyltransferase Esa1 (SMITH *et al*. 1998; Clarke *et al*. 1999), this could explain the essential nature of Spn1. Alternatively, Spn1 depletion may not alter histone acetylation, but rather might impair a different aspect of chromatin structure, such as histone turnover, that can be bypassed by elevated levels of histone acetylation. To begin to distinguish between these possibilities, we performed ChIP-seq to measure the genome-wide levels of histone H4 acetylation. These experiments were done in four conditions: before and after Spn1 depletion in both wild-type *SET2* and *set2Δ* genetic backgrounds. Samples were normalized by adding an equal amount of chromatin from *S. pombe* to each sample prior to immunoprecipitation (see Methods). To assay the levels of acetylation, we used an antiserum that targets histone H4 acetylated at lysines 5, 8, 12, and 16 (Methods), and H4 acetylation levels were compared to total histone H4 levels. Each condition was performed in triplicate and the depletion of Spn1 was verified for each culture by Western blot analysis (Supplementary Figure 1A). Our analysis showed that we had highly reproducible results for most replicates (Supplementary Figure 1B, 1C).

Our results provided several new pieces of information. First, Spn1 depletion alone caused a shift in the pattern of histone H4 acetylation, as we observed a modestly decreased level of H4 acetylation over 5’ regions and a modestly increased level of H4 acetylation over gene bodies after Spn1 depletion (Figure 2A). These changes were most easily observable over long genes (Figure 2A, 2E). These results are consistent with an earlier study that showed an increase in H4 acetylation at two target genes after knockdown of the Spn1 homolog in mammalian cells (Yoh *et al*. 2008). Second, in a *set2Δ* background, both before and after Spn1 depletion, we observed a greatly increased level of H4 acetylation in agreement with the previous studies cited earlier (Figure 2B, 2C, 2E). Third, in the *set2Δ* background, Spn1 depletion still caused a modest increase in H4 acetylation levels (Figure 2D, 2E). This demonstrated that the increased level of histone acetylation upon Spn1 depletion was not merely a consequence of the reduced recruitment of Set2 that results from Spn1 depletion (Reim *et al*. 2020). Together, these results show that Spn1 controls the pattern of histone acetylation across the genome. Furthermore, the results support a model in which bypass of Spn1 depletion by *set2Δ* correlates with hyper-elevated histone acetylation levels.

**Figure 2.**
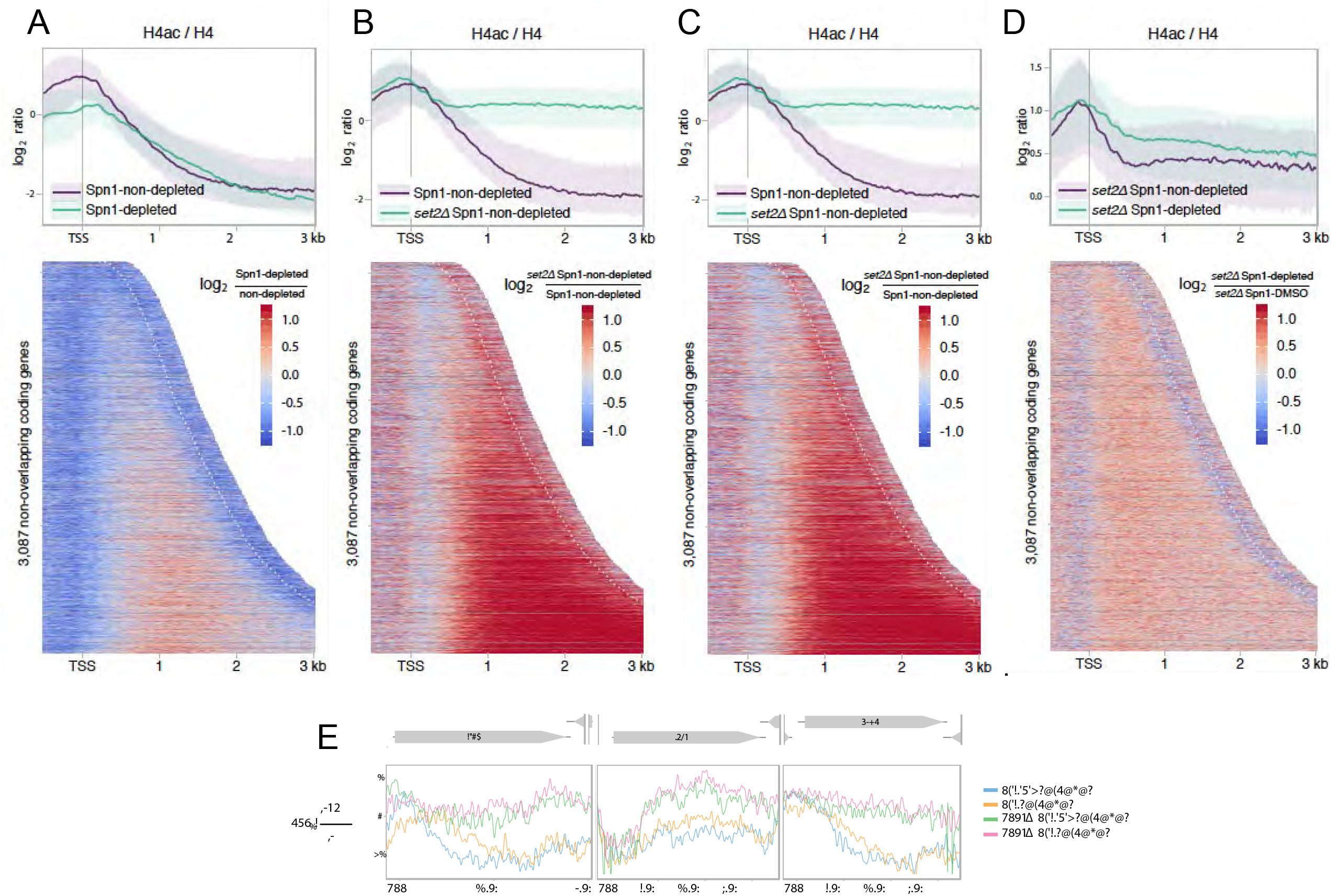
ChIP-seq analysis of histone H4 acetylation. (A) Comparison of histone H4 acetylation levels in strains with and without Spn1 depletion. Top: metagene plots showing the average level of histone H4 acetylation normalized to total levels of histone H4 for 3,087 non-overlapping coding genes in Spn1 non-depleted and depleted conditions. The solid line and the shading are the median and interquartile range of each sample. Bottom: a heatmap showing the same set of genes aligned by the transcription start site (TSS) and arranged by transcript length. The white dotted line on the right represents the position of the cleavage and polyadenylation site. (B) As in panel A except comparing Spn1 non-depleted to *set2Δ* Spn1 non-depleted. (C) As in A except comparing Spn1 non-depleted to *set2Δ* Spn1-depleted. (D) As in A except comparing *set2Δ* Spn1 non-depleted to *set2Δ* Spn1-depleted. (E) Single gene examples of ChIP-seq analysis. The H4 acetylation levels normalized to total H4 levels are shown for three genes, *MYO5, EGT2* and *NHA1*. These three genes showed notable differences in H4ac between the Spn1 depleted and non-depleted conditions. They do not have significant changes in mRNA levels after Spn1 depletion (Reim *et al*. 2020).

In contrast to changes in histone acetylation, we observed that Spn1 depletion had little effect on the level or pattern of total histone H4 levels associated with chromatin. However, Spn1 depletion did cause a decreased level of chromatin-associated histone H4 over the 5’ ends of a small set of genes (Supplementary Figure 1D). These genes tended to be highly expressed, similar to our previous observations for histone H3 (Reim *et al*. 2020) (Supplementary Figure 1E). The results of our ChIP-seq experiments demonstrated a correlation between the elevated levels of histone acetylation in a *set2Δ* background and bypass of Spn1. To test whether histone acetylation is required for Spn1 bypass, we constructed strains to ask whether Gcn5, an H3 histone acetyltransferase (HAT), or Esa1, a histone H4 HAT, were required for suppression by *set2Δ*. These two HATs were chosen to be tested as both have been shown to be recruited to coding regions (Govind *et al*. 2007; Ginsburg *et al*. 2009). As *GCN5* is not essential for viability we tested a *set2Δ gcn5Δ* double mutant. However, *ESA1* is essential for viability, so we tested a temperature-sensitive allele, *esa1-L254P*, which causes reduced HAT activity at increased temperatures (Clarke *et al*. 1999). Our results demonstrated that both Gcn5 and Esa1 are required for suppression by *set2Δ* (Figure 3A, 3B). These results strongly support the model that increased histone acetylation is required for Spn1 bypass by *set2Δ*.

**Figure 3.**
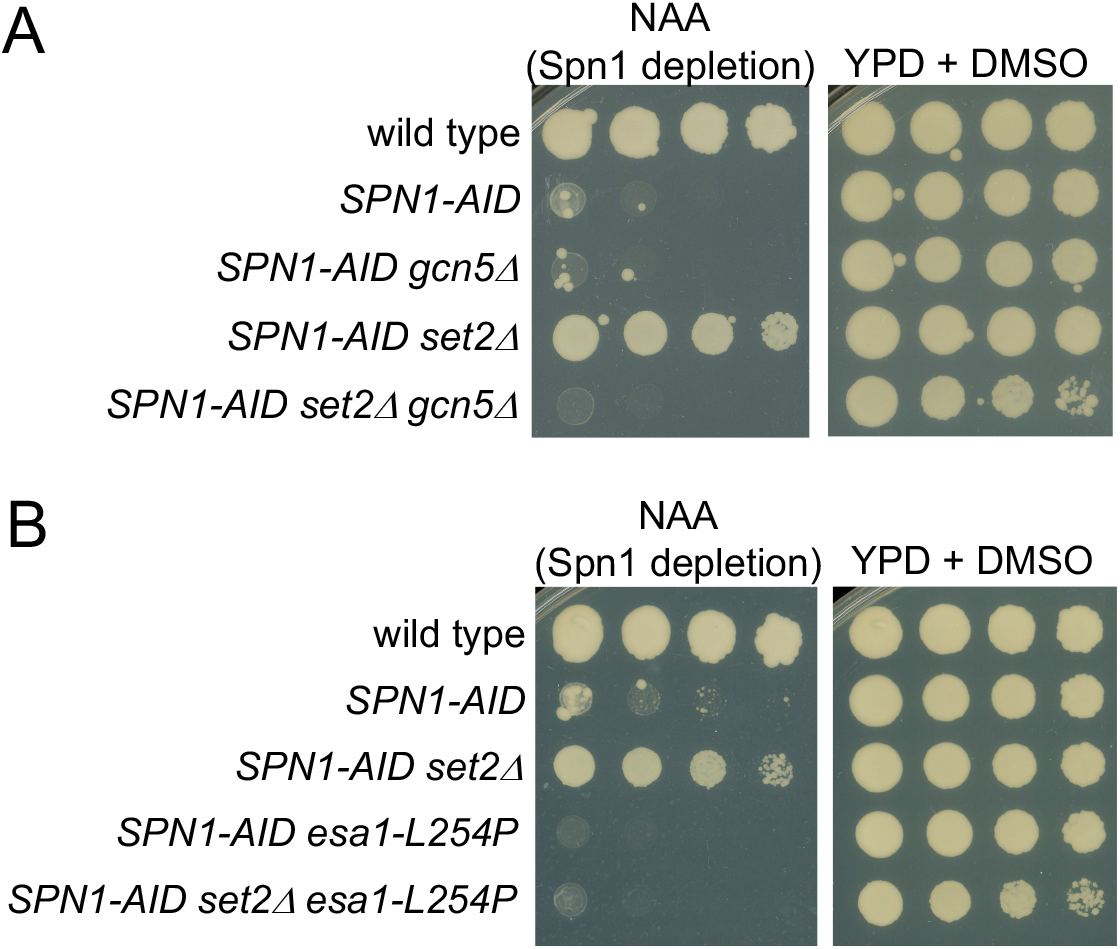
Spn1 bypass by *set2Δ* requires histone acetylation. (A) Dilution spot tests to examine whether Gcn5 is required for Spn1 bypass by *set2Δ*. Plates are shown after three days at 30°C. (B) Dilution spot tests to examine whether Esa1 is required for Spn1 bypass by *set2Δ*. Plates are shown after two days of incubation at 34°C.

### Evidence that Spn1 bypass can occur by multiple mechanisms

Given that Gcn5 is strongly required for *set2Δ* to bypass Spn1, we asked whether Gcn5 is also required for other suppressor mutations to bypass Spn1. To do this, we combined *gcn5Δ* with four other strong suppressors (*rco1Δ, rtt109Δ, chd1Δ*, and *pob3-E154K)* and tested the ability of these strains to survive Spn1 depletion. As expected, *gcn5Δ* abolished suppression by *rco1Δ*, as both Set2 and Rco1 are required for Rpd3S function (Figure 4). In contrast, *gcn5Δ* had little or no effect on suppression by *rtt109Δ, chd1Δ*, or *pob3-E154K* (Figure 4), showing that this latter group of suppressors does not strongly require Gcn5-mediated histone acetylation to bypass Spn1. These results suggest that Spn1 bypass by *set2Δ* and rco1*Δ* occurs by elevated histone acetylation levels, whereas Spn1 bypass by the other suppressors tested, *rtt109Δ, chd1Δ*, and *pob3-E154K*, occurs by one or more distinct mechanisms that are not dependent on Gcn5-dependent histone acetylation.

**Figure 4.**
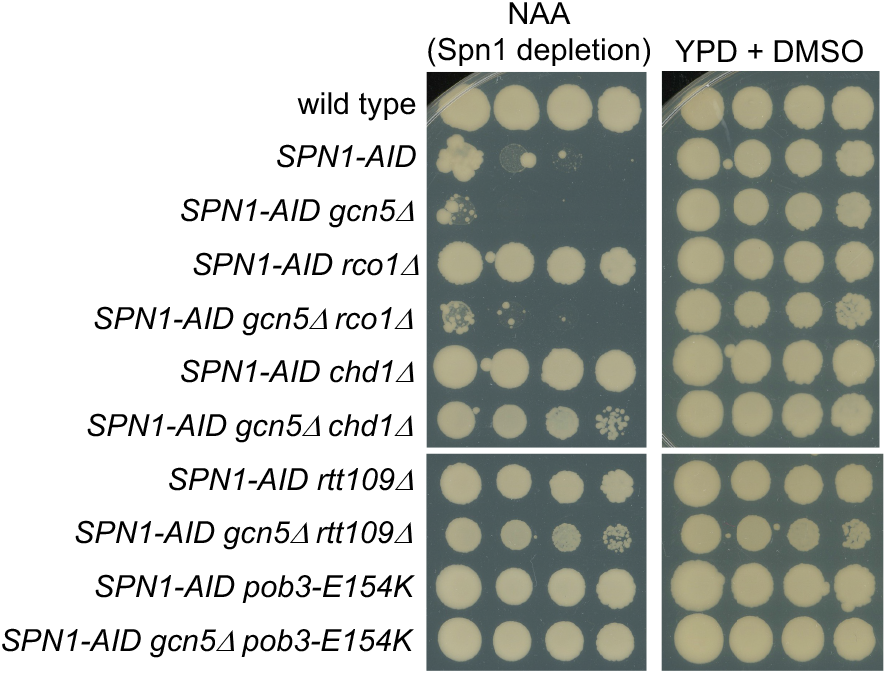
Epistasis tests between Spn1 bypass suppressors and *gcn5Δ*. The plates shown were incubated at 30°C for three days.

### Analysis of Spn1 bypass by *rtt109Δ*

Bypass suppression of Spn1 by *rtt109Δ* was intriguing, as Rtt109 is a histone acetyltransferase that targets multiple sites in histones H3 and H4, several of which overlap the sites targeted by Gcn5 (Schneider *et al*. 2006; Driscoll *et al*. 2007; Han *et al*. 2007; Tsubota *et al*. 2007; Fillingham *et al*. 2008; Burgess *et al*. 2010; Albaugh *et al*. 2011; Abshiru *et al*. 2013; Kuo *et al*. 2015). Therefore, while loss of one HAT, Gcn5, has a negative effect on Spn1 bypass, loss of a different HAT, Rtt109, causes Spn1 bypass. In an attempt to understand how decreased acetylation bypasses Spn1 and how two HATs with overlapping targets cause opposite effects, we investigated the mechanism by which *rtt109Δ* allows Spn1 bypass.

First, as there are multiple H3 HATs in yeast, we tested whether suppression was specific to loss of Rtt109 by directly comparing loss of three different H3 acetyltransferases: Rtt109, Gcn5, and Sas3. Our results showed that only *rtt109Δ* allows Spn1 bypass (Figure 5A). Second, to verify that bypass by *rtt109Δ* is caused by loss of Rtt109 acetylation activity, we tested whether *rtt109-D89A*, which encodes a catalytically dead protein (Han *et al*. 2007), allowed Spn1 bypass. Our results showed that loss of Rtt109 catalytic activity is equivalent to an *rtt109Δ* mutation with respect to Spn1 bypass (Figure 5B). Thus, among the histone H3 HATs studied, Spn1 bypass suppression is specific to loss of Rtt109 acetyltransferase activity.

**Figure 5.**
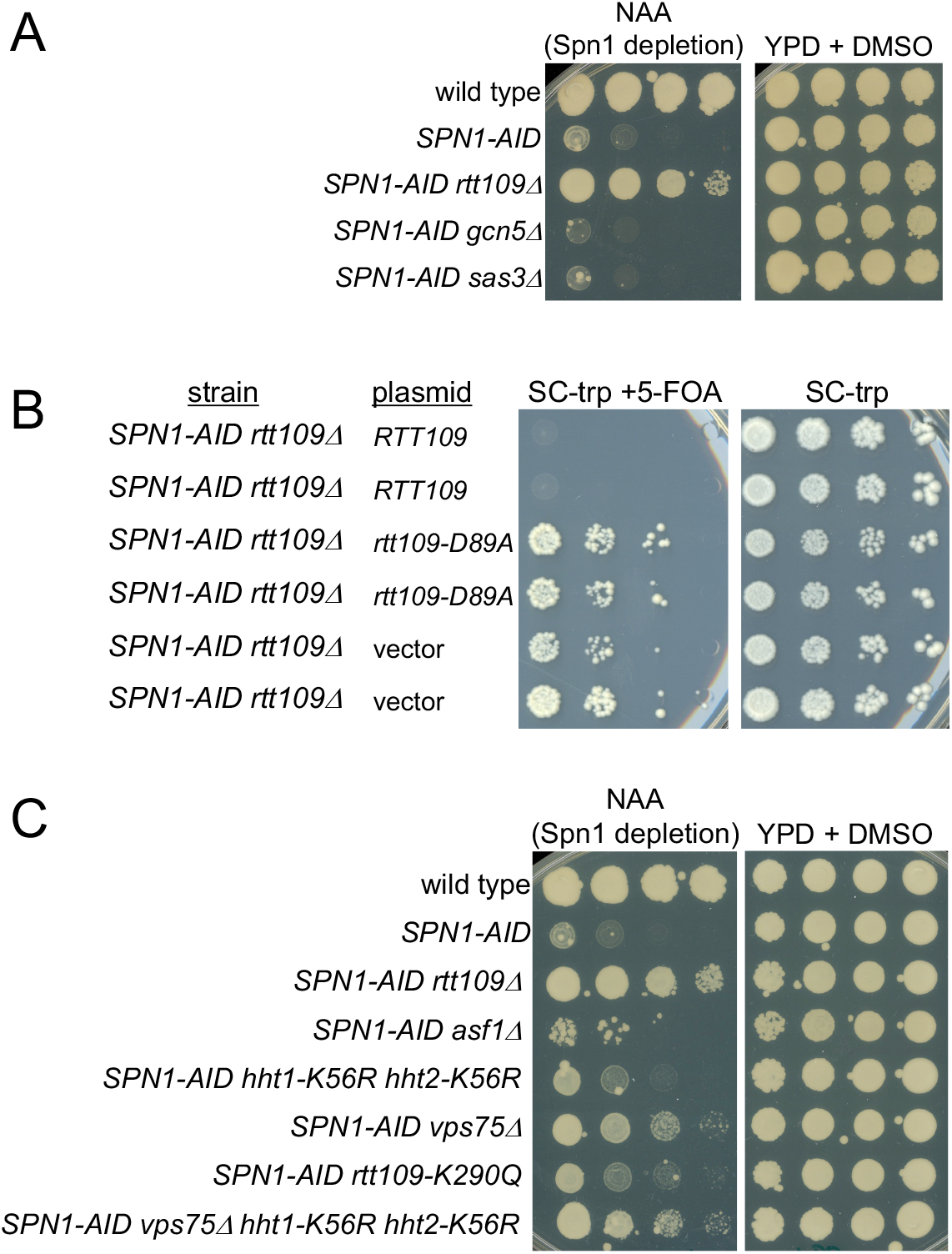
Genetic analysis to understand *rtt109Δ* bypass of Spn1. (A) Dilution spot tests to examine the loss of three different histone acetyltransferases for Spn1 bypass. Plates are shown after three days at 30°C. (B) Dilution spot tests of an *rtt109* catalytically dead mutant. Plates are shown after three days at 30°C. Vector refers to the same plasmid without an *RTT109* gene. (C) Dilution spot tests to analyze mutants that impair different aspects of Rtt109 activity. Plates are shown after three days at 30°C.

Although Rtt109 has multiple acetylation targets, it is the only HAT that acetylates H3K56 (Driscoll *et al*. 2007; Han *et al*. 2007), a histone modification that has been implicated in controlling genome stability (Gershon and Kupiec 2021). Given this unique role for Rtt109, it seemed likely that loss of H3K56 acetylation (H3K56Ac) would be responsible for Spn1 bypass. To test this possibility, we took advantage of previous work that showed that Rtt109 functions in combination with two histone chaperones, Asf1 and Vps75, to help specify the Rtt109 targets. The interaction with Asf1, but not with Vps75, is absolutely required for Rtt109-mediated H3K56 acetylation (Recht *et al*. 2006; Fillingham *et al*. 2008; Zhang *et al*. 2018). When we tested for Spn1 bypass in an *asf1Δ* mutant, our results showed that there was no detectable effect of *asf1Δ* (Figure 5C). To test loss of H3K56 acetylation by an alternative approach, we constructed strains that contained mutations in both copies of the histone H3 genes (*HHT1* and *HHT2*) that mimicked the unacetylated state (H3K56R) (Mccullough *et al*. 2019). In this case, we observed very modest Spn1 bypass, far weaker than was caused by *rtt109Δ* (Figure 5C).

Taken together, these showed that loss of H3K56Ac alone was not sufficient for Spn1 bypass. To test if bypass occurs by loss of acetylation at other Rtt109 histone target sites, we tested two mutations, *vps75Δ* (Selth and Svejstrup 2007; Tsubota *et al*. 2007; Fillingham *et al*. 2008) and *rtt109-Q290K* (Radovani *et al*. 2013), that primarily decrease Rtt109 acetylation at sites other than K56. Our results showed that both *vps75Δ* and *rtt109-Q290K* conferred very weak Spn1 bypass, similar to the H3K56R changes (Figure 5C). When we tested a combination of mutations, H3K56R with *vps75Δ*, that would impair Rtt109 acetylation at most or all of its known histone targets, we observed slightly greater, but still quite modest Spn1 bypass (Figure 5C). In summary, by several approaches to independently impair Rtt109 acetylation at its histone targets, the mutants were unable to recapitulate the level of Spn1 bypass observed by either *rtt109Δ* or a mutation that impairs Rtt109 catalytic activity. These results suggest the existence of additional Rtt109 targets that are independent of Vps75 and Asf1.

## Discussion

Spn1 is a highly conserved and essential transcription factor that functions as a histone chaperone and that plays critical roles in transcription and chromatin structure. We have now shown that the inviability caused by the loss of Spn1 can be bypassed in *S. cerevisiae* by single mutations in eight different genes that play distinct roles in transcription and chromatin structure. Our results suggest that bypass can occur by distinct mechanisms. In the clearest case, bypass of Spn1 by *set2Δ*, we showed that bypass required increased levels of histone acetylation. In contrast, increased histone acetylation was not required in other cases, such as Spn1 bypass by either *chd1Δ* or *pob3* mutations. Furthermore, bypass of Spn1 by *rtt109Δ* required loss of the Rtt109 acetyltransferase activity, rather than elevation of acetylation. The bypass of Spn1 by different mechanisms suggests that Spn1 may have multiple functions *in vivo*, that the combination of these functions results in the essential nature of Spn1, and that the suppression of a subset of these functions can bypass the need for Spn1 for viability.

Bypass of Spn1 by elevated histone acetylation levels suggests that Spn1 normally promotes DNA accessibility, and in its absence, chromatin remains in a repressed state that prevents transcription, and possibly other chromatin-related processes, from occurring normally. There are several possible reasons why increased histone acetylation would allow bypass. First, increased acetylation of histone N-terminal tails has been shown to lead to reduced electrostatic interactions of the histone tails with DNA, loosening chromatin and making the DNA more accessible to transcription factors (Protacio *et al*. 2000). In addition, histone acetylation recruits bromodomain-containing proteins, including nucleosome remodelers such as Swi/Snf (Hassan *et al*. 2002) and RSC (Carey *et al*. 2006; Ginsburg *et al*. 2009; Mas *et al*. 2009), and there is evidence that histone acetylation also facilitates histone turnover (Venkatesh *et al*. 2012). There is evidence that histone acetylation levels affect the recruitment of histone chaperones, as one recent study showed that reduced histone acetylation decreased the recruitment of FACT (Pathak *et al*. 2018). However, that same study showed that elevated histone acetylation levels did not increase the recruitment of FACT. Therefore, increased association of FACT seems unlikely to be a contributing factor.

In contrast to *set2Δ*, Spn1 bypass by *chd1Δ* may be caused by increased association of the FACT complex. One recent study showed that in *chd1Δ* mutants there was an increased level of stable association of FACT with chromatin, suggesting a faster “on” rate in the absence of Chd1 (Jeronimo *et al*. 2021). This result is consistent with recent structural studies that showed that the binding of FACT and Chd1 to a nucleosome was mutually exclusive (Farnung *et al*. 2021).

Thus, removing Chd1 by mutation might provide FACT with an increased opportunity to associate with transcribed chromatin. By this model, it is possible that some of the *spt16* and *pob3* bypass suppressors isolated in this study would increase FACT activity. This model also fits with previous studies that showed that *chd1Δ* suppresses a distinct class of *spt16* and *pob3* temperature-sensitive mutations (Simic *et al*. 2003; Biswas *et al*. 2007). In those cases, the decreased FACT activity caused by those *spt16* and *pob3* mutations might be compensated by an increased association of FACT with chromatin. Other changes that have been shown to occur in *chd1Δ* mutants are less likely to contribute to Spn1 bypass, including increased levels of histone acetylation and altered localization of some histone modifications (Radman-livaja *et al*. 2012; Smolle *et al*. 2012; Lee *et al*. 2017), given the Gcn5-independence of *chd1Δ* bypass.

Although our experiments did not elucidate the mechanism by which *rtt109Δ* bypasses Spn1, they did rule out specific possibilities and point toward future areas for investigation of Rtt109. First, by multiple genetic tests, we have eliminated the possibility that Spn1 bypass occurs solely via loss of H3K56 acetylation by Rtt109. In addition, our results showed that loss of acetylation at all of the known Rtt109 histone targets fails to recapitulate Spn1 bypass by *rtt109Δ*. Since loss of Rtt109 catalytic activity is sufficient for Spn1 bypass, our results suggest that Spn1 bypass by *rtt109Δ* requires loss of acetylation at novel Rtt109 targets that have not yet been identified. Such targets could be sites in histones that are independent of Asf1 and Vps75, or possibly, non-histone targets.

Several recent structural studies have provided important new insights into the structure of the RNAPII eukaryotic transcription elongation complex and its interactions with nucleosomes (Ehara *et al*. 2017; Farnung *et al*. 2018; VOS *et al*. 2018a; Vos *et al*. 2018b; EHARA *et al*. 2019; VOS *et al*. 2020; Farnung *et al*. 2021). Although these structures have not yet revealed the location and detailed interactions of Spn1 within this complex, multiple studies have shown that Spn1 is part of the elongation complex (Krogan *et al*. 2002; Lindstrom *et al*. 2003; Zhang *et al*. 2008; Diebold *et al*. 2010; Mayer *et al*. 2010; Mcdonald *et al*. 2010; Pujari *et al*. 2010; Li *et al*. 2018; Reim *et al*. 2020; Cermakova *et al*. 2021; VIKTOROVSKAYA *et al*. 2021), and recent evidence suggests that, in human cells, Spn1 serves as a hub for the elongation complex (Cermakova *et al*. 2021). Our results have demonstrated that, in the presence of bypass suppressors, yeast can survive, and in some cases grow at close to wild-type rates, in the absence of Spn1. It will be of great interest to learn how Spn1 localizes within the transcription elongation complex and to understand whether the requirement for Spn1 can be bypassed in larger eukaryotes as well as in yeast.

### Data availability

All strains and plasmids are available upon request. Sequencing data from this study is available at the NCBI Gene Expression Omnibus (GEO) with the accession number GSE202590. The code used for the ChIP-seq analysis is available at https://github.com/orgs/winston-lab/repositories.

## Acknowledgments

We thank Catherine Miller, Natalia Reim, and Olga Viktorovskaya for helpful comments on the manuscript. We thank Lorraine Pillus, Alain Verreault, Sharon Dent, Andrew Salinger, Tim Formosa, Laura McCullough, Karen Arndt, Jeffrey Fillingham, Grant Hartzog, Stephen Buratowski, Laurie Stargell, and Zhiguo Zhang for strains, plasmids, and antibodies.

